# Linking Covariates to State Durations in Dynamic Brain Networks via Hidden Semi-Markov Models

**DOI:** 10.64898/2026.11.06.764975

**Authors:** Heather M. Shappell, Martin A. Lindquist

## Abstract

Hidden semi-Markov models (HSMMs) provide a flexible framework for studying dynamic functional connectivity because they represent brain activity as transitions between latent network states with explicit dwell-time distributions. Although dwell times are often associated with behavioral and demographic variables, these associations are typically assessed only after model fitting using indirect group comparisons, limiting formal covariate modeling and the simultaneous inclusion of multiple predictors. In this work, we extend the HSMM framework by introducing covariate-dependent dwell-time distributions through a regression-based formulation, allowing expected state duration to vary with subject-level covariates and enabling direct in-ference on how these factors relate to state persistence. Model parameters are estimated with an expectation–maximization algorithm, and uncertainty in the dwell-time regression coefficients is quantified using a subject-level nonparametric bootstrap. In simulations, the proposed model accurately recovered covariate effects under both shifted-Poisson and Gamma dwell-time specifications, with low bias and variability. We applied the method to resting-state fMRI data from 1,000 UK Biobank participants and found that age and sex were associated with dwell times in a subset of brain states. Together, these results show that modeling dwell-time behavior directly provides a more flexible framework for studying individual differences in brain dynamics, revealing not only which states occur, but also how long individuals remain in them.

## 1. Introduction

Interest in studying brain networks, in which the brain is conceptualized as a system of interacting regions (nodes) that give rise to complex behaviors, has grown tremendously over the past decade. Networks have become a popular approach for characterizing both the physiological connections (structural networks) and the coupling of dynamic brain activity (functional networks) linking different areas of the brain. Within this broader shift, scientists have begun investigating how networks behave in healthy brains and how they are altered in neurological and psychiatric disorders (Stam (2014)).

The study of functional connectivity (FC), or the undirected association between two or more fMRI time series, has come to the forefront of research efforts in the field of neuroimaging. Such studies have revealed important information about the functional relationships among brain regions and local networks and have provided significant insights into the organization of functional communication in the brain. Brain networks can be created by performing an FC analysis, where the strength of the relationship between nodes is assessed using the functional magnetic resonance imaging (fMRI) time series associated with each node. The nodes can consist of individual voxels, pre-specified regions of interest (ROIs), or sets of regions estimated using independent component analysis (ICA). The relationship between nodes can be quantified using a variety of metrics, such as Pearson and partial correlation.

To date, most of the brain connectivity analyses performed estimate a single aggregate brain network that does not change during a specific experimental fMRI scanning session (Donofry et al. (2020)). While convenient, this oversimplifies the functional relationship between brain regions. Empirical studies indicate that brain networks are dynamic and changing, even over brief time periods (Chang and Glover (2010); Hutchison et al. (2013)) thereby providing insight into behavioral shifts and adaptive processes in both normal and abnormal brain function. Thus, there is a need to estimate the timing of the transitions between states, as well as the structure of the states themselves. Several approaches for assessing time-varying connectivity have been suggested. These include using sliding window correlations (Allen et al. (2014)), change point models (Cribben et al. (2012, 2013); Xu and Lindquist (2015)), wavelet transform coherence (Chang and Glover (2010)), and time series models (Lindquist et al. (2014)).

In recent years, Hidden Markov models (HMMs) have also proven to be a useful modeling approach for assessing FC (Vidaurre et al. (2017)). In this context, the time series from each brain region is assumed to arise from a sequence of a fixed number of hidden brain states. Each state is characterized by a multivariate Gaussian distribution, which is parameterized by its mean and covariance. Of particular interest is the unique connectivity structure of each state, encapsulated within the covariance matrix. An HMM approach to dynamic FC analysis is appealing for several reasons. First, the framework can handle large amounts of data to infer the network states, along with other model parameters such as the probabilities of transitioning between states. Moreover, these states and parameters can be estimated in a relatively computationally inexpensive manner. Second, while network states are estimated at the group level, individuality is respected in that information about when a state becomes active is subject-specific. Therefore, the model still yields subject-specific estimates of an individual’s state sequence and the amount of time spent in each state. Third, the model is able to capture quick changes in brain connectivity, unlike the sliding-window-based approach.

Unfortunately, despite these strengths, the HMM framework relies on a potentially unrealistic assumption, which is that the dwell-time, or the number of consecutive time points spent in a specific state, is geometrically distributed. A property of the geometric distribution is that the probability decreases as the dwell-time increases. Therefore, more weight is placed on shorter consecutive time points in a given state. This may not be appropriate in the context of functional connectivity analysis, as it suggests that subjects are switching states very often, as opposed to potentially spending longer periods of time in the same state. For example, in some task-based fMRI experiments, participants are expected to spend several minutes in a particular brain state. It also suggests that the dwell times for all states follow a very similar pattern, which may not be the case in practice.

To address this limitation, in our prior work, we proposed explicitly modeling and estimating the brain state dwell-time distributions via hidden semi-Markov models (HSMMs) (Shappell et al. (2019)). HSMMs differ in that the dwell distributions are explicitly built into the likelihood function. One may specify the distributions to take a nonparametric form or a parametric form (such as a Poisson or Gamma distribution) and estimate the parameters directly. In that work, the HSMM was used to estimate network states common across subjects in a particular study. Moreover, the parameter estimates we obtain from fitting the HSMM allow us to find each individual subject’s most probable series of network states along with the amount of time spent in each state. The estimated model also provides additional information, such as the probabilities of transitioning from one network state to another and the dwell-time distributions for each state.

However, the HSMM as it is currently formulated does not allow one to directly test for associations between the state dwell-time distributions and various effects, nor does it allow one to test for these associations while controlling for potential confounders. It is possible to test such associations indirectly, in an ad hoc manner, for a single categorical variable and a state dwell-time distribution, but this would need to be done via permutation testing. This approach does not allow for the assessment of continuous covariates or the simultaneous inclusion of multiple effects.

This is a significant limitation of the current model, as dwell times have been shown to provide valuable information about the brain mechanisms underlying various cognitive processes, and they are an important component of studying dynamic properties of brain networks. For example, Chen et al. (2025) found evidence across multi-center datasets that creativity is tied to the capacity to dynamically switch between brain networks supporting spontaneous and controlled cognition. How long an individual dwells in a particular state is closely linked to when and how often they switch between different brain networks. Moreover, Pedersen et al. (2018) showed, using a multilayer model, that brain regions, particularly the lateral frontal and parietal brain areas, transition between different network configurations at a high rate (i.e., they have short dwell times). They found that this network switching rate predicts performance of higher-order cognitive functions including working memory, planning, and reasoning. In other words, efficient brain network switching appears to be an important aspect of optimal brain function.

In our own work, we have found that dwell-time distributions were particularly discriminative across attention-deficit/hyperactivity disorder (ADHD) and typically developing (TD) groups, where children with ADHD had trouble traversing in and out of certain states. Specifically, those with ADHD got ‘stuck’ in an inattentive state characterized by high connectivity between the DMN and task-relevant networks, while they were less likely to enter into or maintain a cognitive control, or on-task attention, state characterized by anticorrelations between the DMN and task-relevant networks (Shappell et al. (2021)).

Therefore, in this work, we propose a covariate-dependent HSMM that allows the dwell-time distribution to depend on subject-level, time-invariant fixed effects. The model retains the ability to estimate network states shared across subjects while allowing state dwell times to depend directly on these covariates. This enables formal estimation and inference for covariate effects on state persistence. The implementation supports shifted-Poisson and Gamma dwell-time specifications.

In the following sections, we present the model, evaluate its performance through simulation, and apply it to UK Biobank data. The results show that the proposed approach accurately recovers dwell-time covariate effects and identifies associations between state dwell times and both sex and age, underscoring the utility of this framework for studying individual differences in brain dynamics.

## 2. Model Formulation

### 2.1. Hidden semi-Markov model for fMRI data

We assume that we observe resting-state fMRI time series data *Y*_1_, …, *Y*_*T*_, where each vector *Y*_*t*_ ∈ ℝ^1×*p*^ contains the BOLD measurements from *p* brain regions at the *t*^*th*^ repetition time (TR). Throughout this paper, we model each *Y*_*t*_ as arising from one of *K* latent brain states. Conditional on the underlying state, the observed BOLD vector follows a multivariate Gaussian distribution,

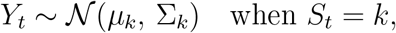

where *µ*_*k*_ ∈ ℝ^1×*p*^ represents the mean activation pattern, and Σ_*k*_ ∈ ℝ^*p*×*p*^ captures the functional connectivity structure among the *p* regions in state *k*. Thus, each state has its own unique mean activation pattern across the *p* brain regions, as well as its own connectivity structure describing the interactions between each pair of brain regions.

The collection of observed BOLD vectors *Y*_1_, …, *Y*_*T*_ is denoted by 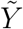, and the unobserved latent brain states at each TR are denoted by *S*_1_, …, *S*_*T*_. We write the full latent state sequence as 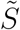, with each *S*_*t*_ ∈ {1, …, *K*}.

### 2.2. Latent State Process

The latent state process is modeled as a semi-Markov chain. Specifically, once a brain state is entered, the process remains in that state for a random dwell-time before transitioning to a new state. Let *s*_*r*_ denote the *r*^*th*^ visited brain state and let *u*_*r*_ denote the dwell-time, defined as the number of con-secutive TRs spent in that state. Let *R* denote the total number of visited states over the observed time series. Transitions between visited states are governed by a conditional transition probability matrix *P*, where

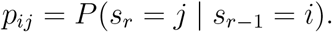

Self-transitions are not permitted, so *p*_*ij*_ = 0, because persistence within a state is represented explicitly through the dwell-time distribution. Unlike a standard hidden Markov model, the dwell time in each state is explicitly modeled through a state-specific distribution *d*_*k*_(*u*) supported on the positive integers.

### 2.3. Complete Data Log-Likelihood

Under this formulation, the complete-data log-likelihood for a single subject can be written as

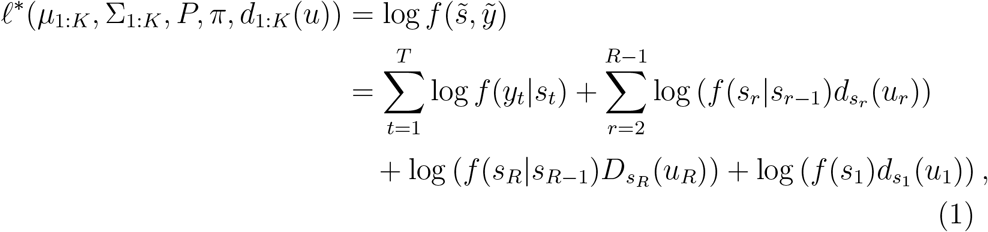

where:

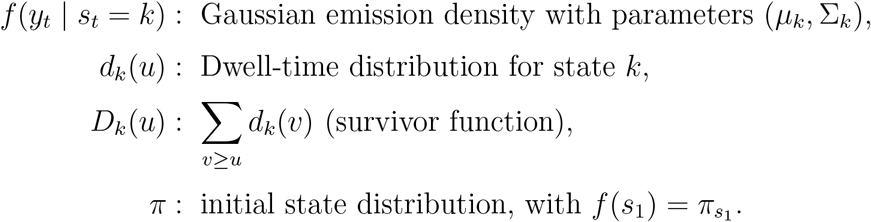

The survivor contribution 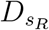 (*u*_*R*_) is used for the final state visit because the observed time series may end before the full duration of that visit is observed. For computation, the dwell-time distribution is evaluated on the finite grid *u* = 1, …, *M*, where *M* = `max_dwell` is the maximum modeled dwell duration. Accordingly, the survivor quantity used in the E-step is computed as

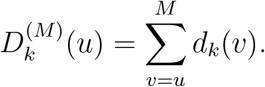

To assess whether the selected value of *M* is sufficiently large, the implementation examines the proportion of the evaluated dwell distribution contained in the final five grid points and flags fitted models with appreciable mass near the upper boundary.

Now suppose data are observed from *N* independent subjects, each contributing an fMRI time series 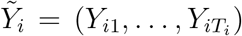 and associated covariate vector *x*_*i*_. Assuming independence across subjects, the full observed-data likelihood is

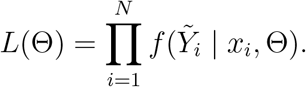

Parameter estimation proceeds by maximizing this likelihood using the Expectation–Maximization (EM) algorithm described in Section 2.5.

If *d*_*k*_(*u*) were geometric, this formulation would reduce to the standard hidden Markov model. However, in dynamic functional connectivity applications, a geometric dwell-time distribution may place unrealistic weight on very short state visits. In contrast, the HSMM estimates dwell-time distributions directly from the data, providing greater flexibility in characterizing the temporal stability of functional brain configurations. More generally, HSMM dwell-time distributions may be specified parametrically or nonparametrically. The implementation considered here supports shifted-Poisson and Gamma dwell-time specifications. In the formulation above, the dwell-time distribution depends only on the state *k* and is therefore assumed to be identical across subjects; that is, *d*_*k*_(*u*) depends only on the state and not on subject-level characteristics. In the following section, we extend this framework to allow dwell-times to depend on subject-level, time-invariant covariates.

### 2.4. Covariate-Dependent Dwell-Time Modeling

We now extend the HSMM framework to allow the dwell-time distribution to depend on subject-level, time-invariant covariates. Let *x*_*i*_ ∈ ℝ^*q*^ denote the design vector for subject *i*, including an intercept and any indicator variables introduced through covariate coding. A single design vector is associated with each subject and applies to all state visits for that subject. Under this extension, the dwell-time specification for state *k* is no longer identical across subjects, but instead depends on *x*_*i*_ through a state-specific regression structure. We write

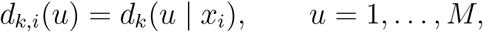

where *d*_*k,i*_(*u*) denotes the dwell-time contribution evaluated at integer duration *u* for subject *i* in state *k*, and *M* is the maximum modeled dwell duration. Let *U*_*ir*_ denote the dwell time associated with the *r*^*th*^ visited state for subject *i*. Under the shifted-Poisson specification, *d*_*k,i*_(*u*) is a probability mass function. Under the Gamma specification, *d*_*k,i*_(*u*) is obtained by evaluating the continuous Gamma density at the integer durations *u* = 1, …, *M*, and is used as an integer-grid approximation to the dwell-time contribution, as described below. Importantly, the emission distribution parameters (*µ*_*k*_, Σ_*k*_) and the transition probability matrix *P* remain common across subjects. Only the dwell-time component is allowed to depend on covariates.

#### 2.4.1. Shifted-Poisson Dwell-Time Regression

One possible parametric specification assumes that the dwell time follows a shifted-Poisson distribution supported on the positive integers. Specifically, let

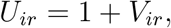

where

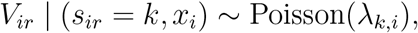

and

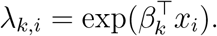

Here, *β*_*k*_ ∈ ℝ^*q*^ is a state-specific regression coefficient vector. The resulting dwell-time probability mass function is

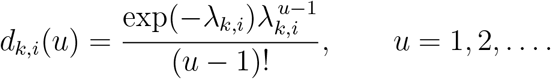

The conditional mean and variance are

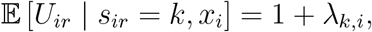

and

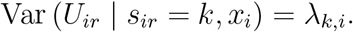

Thus, exp(*β*_*kj*_) represents the multiplicative change in *λ*_*k,i*_ associated with a one-unit increase in covariate *j*, holding the remaining covariates fixed. Because

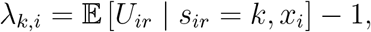

this is a multiplicative effect on the expected dwell time minus one, rather than on the complete expected dwell time.

#### 2.4.2. Gamma Dwell-Time Regression

Alternatively, dwell times may be represented using a Gamma density. The Gamma specification provides additional flexibility in the dispersion and tail behavior of dwell times through a state-specific shape parameter. This can be useful in dynamic functional connectivity analyses, where dwell times may exhibit substantial heterogeneity across visits and subjects.

For subject *i* in state *k*, define the covariate-dependent mean dwell time as

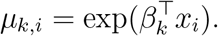

The corresponding Gamma density has state-specific shape parameter *α*_*k*_ *>* 0 and subject-specific scale parameter

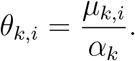

Under this mean–shape parameterization,

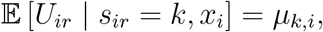

and

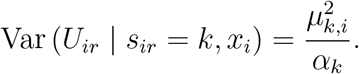

Thus, exp(*β*_*kj*_) represents the multiplicative change in expected dwell time associated with a one-unit increase in covariate *j*, holding the remaining covariates fixed, while *α*_*k*_ controls the relative dispersion of dwell times within state *k*.

Because dwell durations are represented on an integer grid, the implementation evaluates the continuous Gamma density at *u* = 1, …, *M*. Specifically,

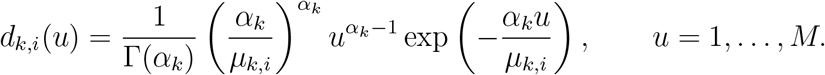

These density values are used directly as the dwell-time contributions in the E-step and are not converted to probabilities over unit-width intervals or renormalized over the finite grid.

### 2.5. Parameter Estimation via EM Algorithm

Model parameters are estimated using the Expectation–Maximization (EM) algorithm (Moon (1996)). The EM algorithm alternates between computing the expected complete-data log-likelihood given the observed BOLD time series and the current parameter estimates (E-step), and maximizing this expectation with respect to the model parameters (M-step). Under the Gamma specification, references to the likelihood correspond to the integergrid density approximation described above.

Let Θ denote the full set of model parameters. Under the shifted-Poisson specification,

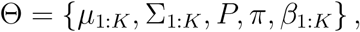

whereas the Gamma specification additionally includes the state-specific shape parameters:

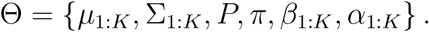

#### 2.5.1. E-Step

Let 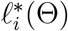 denote the complete-data log-likelihood contribution for subject *i*, and let

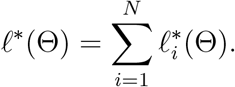

In the E-step, we compute

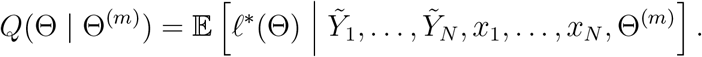

This step requires the following posterior quantities:

1. The marginal posterior probability that subject *i* occupies state *k* at time *t*:

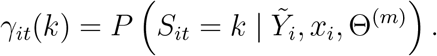
2. The pooled posterior expected number of transitions from visited state *j* to visited state *k*:

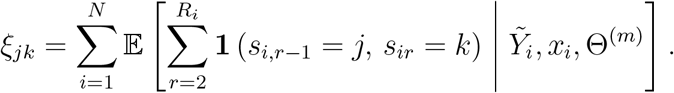
3. The posterior expected number of visits by subject *i* to state *k* having dwell time *u*:

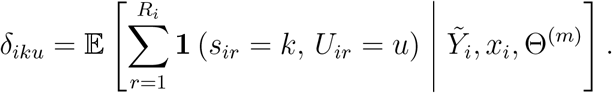

Here, **1**(·) denotes the indicator function, *s*_*ir*_ denotes the *r*^*th*^ visited state for subject *i, U*_*ir*_ is its corresponding dwell time, and *R*_*i*_ is the total number of visited states for subject *i*. For computational efficiency, posterior expected dwell counts are aggregated across subjects having identical covariate design vectors before the dwell-time parameters are updated.

#### 2.5.2. M-Step

In the M-step, we maximize *Q*(Θ | Θ^(*m*)^) with respect to each group of parameters.

##### Emission Parameters

The Gaussian emission parameters are updated by pooling posterior-weighted observations across all subjects and time points:

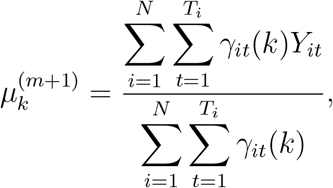

and

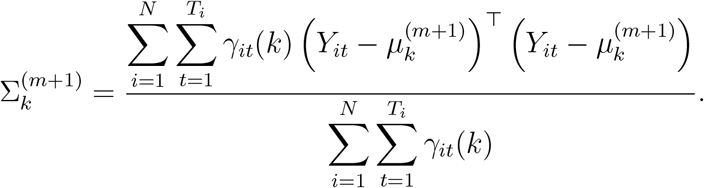

##### Transition Probabilities

Transition probabilities are updated using the pooled expected numbers of transitions between visited states:

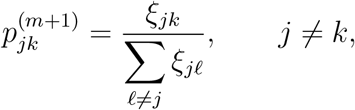

with

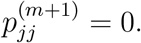

##### Initial Distribution

The initial state probabilities are updated by averaging the posterior state probabilities at the first time point across subjects:

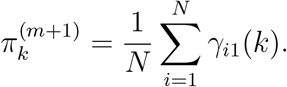

##### Shifted-Poisson Dwell-Time Regression

Under the shifted-Poisson specification,

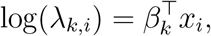

so that

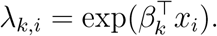

Ignoring terms that do not depend on *β*_*k*_, the expected complete-data log-likelihood contribution for state *k* is

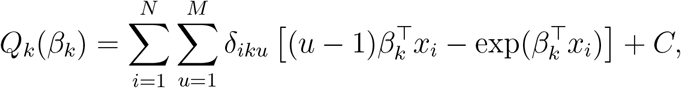

where *C* does not depend on *β*_*k*_.

For computational efficiency, posterior expected dwell counts are aggregated across subjects having identical covariate design vectors. The regression coefficients are then estimated separately for each state by minimizing the negative expected complete-data log-likelihood using bounded L-BFGS-B optimization. This objective is equivalent to a weighted Poisson regression for the shifted duration *u* − 1.

##### Gamma Dwell-Time Regression

Under the Gamma specification,

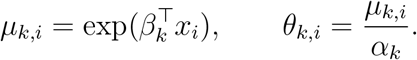

The expected complete-data log-likelihood contribution for state *k* is

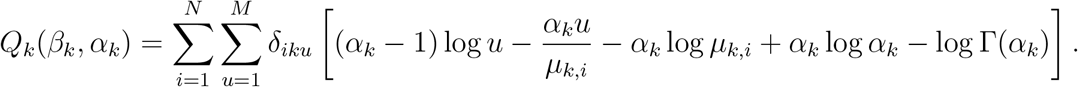

For each state, the posterior expected dwell counts are first aggregated across subjects having identical covariate design vectors. Let *g* index the resulting covariate groups, and define

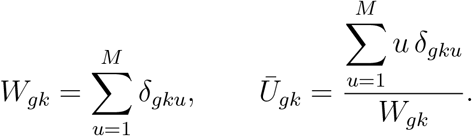

The regression coefficients *β*_*k*_ are updated using a weighted Gamma generalized linear model with a log link, with 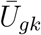 as the response and *W*_*gk*_ as the weight. Under the mean–shape parameterization, the maximizing value of *β*_*k*_ does not depend on *α*_*k*_. Conditional on the updated regression coefficients, the state-specific shape parameter *α*_*k*_ is estimated through one-dimensional numerical optimization over log(*α*_*k*_).

The E- and M-steps are iterated until the absolute change in the total observed-data log-likelihood between successive iterations satisfies

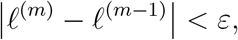

where *ε* is a prespecified convergence tolerance. Convergence is assessed beginning with the third EM iteration. The implementation also flags decreases in the observed-data log-likelihood and declares convergence only when any decrease is within a small numerical tolerance.

### 2.6. Statistical Inference

The EM algorithm described above yields estimates of the model parameters, including the state-specific regression coefficients *β*_*k*_ governing the covariate-dependent dwell-time distributions. However, the conditional optimization steps in the M-step do not directly provide standard errors that account for uncertainty in the latent state sequence and the other estimated model parameters. In particular, the regression updates treat the posterior expected dwell counts *δ*_*iku*_ obtained from the E-step as fixed. Consequently, uncertainty estimates based only on a single M-step update would not reflect the variability associated with estimating the complete HSMM. Because inference on population-level covariate effects is a primary objective of this framework, standard errors for the regression coefficients are estimated using a subject-level nonparametric bootstrap.

The current implementation performs bootstrap inference for the dwell-time regression coefficients *β*_*k*_. Under the Gamma specification, the state-specific shape parameters are re-estimated within each bootstrap replicate, but bootstrap standard errors and hypothesis tests are not currently reported for those parameters.

#### 2.6.1. Bootstrap Procedure

Suppose data are observed from *N* independent subjects, each contributing an fMRI time series and associated covariate design vector *x*_*i*_. Let 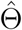 denote the full set of parameter estimates obtained by fitting the covariate-dependent HSMM to the original dataset.

The bootstrap procedure proceeds as follows:

1. For *b* = 1, …, *B*:
  a. Sample *N* subjects with replacement from the original dataset.
  b. For each selected subject, retain the entire observed time series and the corresponding row of the original covariate design matrix. This preserves the original factor coding and any transformations used to construct the design matrix.
  c. Fit the full EM algorithm to the resampled dataset, obtaining bootstrap parameter estimates 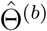, including 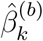 for each state *k*. By default, the parameter estimates from the original full-data fit are used as starting values, but the model is fully refitted within each bootstrap replicate.
  d. Align the fitted states from the bootstrap replicate to the states from the original full-data fit.

Because latent-state models are invariant to permutations of the state labels, state indices may differ across bootstrap replicates. To ensure comparability, the bootstrap states are aligned to the original fitted states using a globally optimal one-to-one assignment. For original state *j* and bootstrap state *k*, the alignment cost is

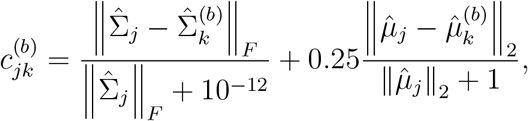

where ∥·∥_*F*_ denotes the Frobenius norm and ∥·∥_2_ denotes the Euclidean norm. The assignment minimizing the total alignment cost across states is obtained using the Hungarian algorithm (Kuhn (1955)). Thus, state alignment is based primarily on similarity between the emission covariance matrices, with a smaller contribution from similarity between the emission mean vectors.

Bootstrap replicates that result in errors, rank-deficient resampled design matrices, or nonfinite regression coefficient estimates are excluded from inference. By default, replicates that do not satisfy the EM convergence criterion are also excluded. Let B_use_ denote the set of usable bootstrap replicates and let

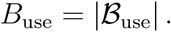

The numbers of requested, successfully fitted, converged, and usable boot-strap replicates are recorded and reported.

For a given state *k* and covariate coefficient *β*_*k,ℓ*_, define the mean boot-strap estimate across usable replicates as

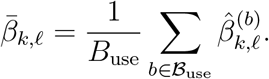

The bootstrap standard error is computed as the empirical standard deviation across the usable bootstrap replicates:

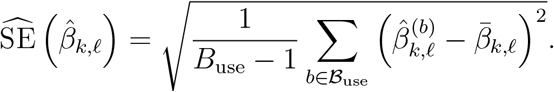

#### 2.6.2. Hypothesis Testing

Inference on covariate effects is conducted separately for each state and regression coefficient. For example, to assess whether age is associated with dwell time in state *k*, we test

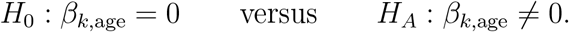

A Wald statistic is calculated as

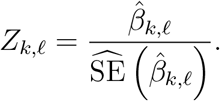

Two-sided *p*-values are obtained using the standard normal reference distribution:

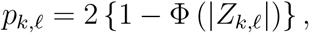

where Φ(·) denotes the standard normal cumulative distribution function.

The implementation reports a 95% normal-approximation confidence interval,

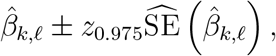

as well as a 95% percentile bootstrap interval,

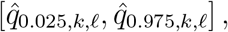

where 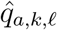 is the empirical *a*-quantile of 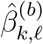 across the usable bootstrap replicates. By resampling entire subjects and fully refitting the model, the boot-strap incorporates variability associated with subject sampling, latent-state estimation, and re-estimation of the other model parameters. It therefore provides a more comprehensive assessment of uncertainty in the dwell-time regression coefficients than an uncertainty estimate conditional on a single M-step update.

## 3. Simulation Study

We conducted a simulation study to evaluate the ability of the proposed covariate-dependent HSMM to recover associations between subject-level covariates and state dwell-time distributions. In particular, we assessed the accuracy with which the model recovered the state-specific regression coefficients governing dwell times under both shifted-Poisson and Gamma dwell-time formulations.

Simulations were performed under two data-generating scenarios. In the first scenario, dwell times followed a shifted-Poisson distribution, while in the second they were generated from a Gamma distribution. In both cases, subject-specific latent state sequences were first generated from an HSMM with covariate-dependent dwell times, after which multivariate fMRI-like time series were simulated conditional on the latent state sequence.

### 3.1. Simulation Design

For each simulation scenario, we generated 100 independent datasets. Each dataset consisted of *N* = 400 subjects, with *T* = 500 time points per subject. The latent process contained *K* = 4 states. Initial state probabilities were set to

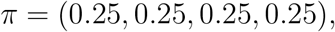

and the transition probability matrix governing movement between visited states was

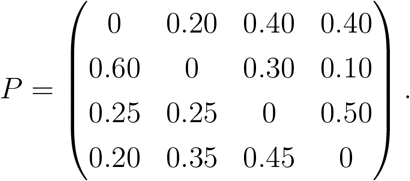

Thus, self-transitions were not allowed in the transition matrix because persistence within a state was represented explicitly through the dwell-time distribution.

Each subject was assigned one binary covariate and one continuous covariate. The binary covariate was generated so that, within each simulated dataset, 200 subjects had value 0 and 200 subjects had value 1. The continuous covariate was generated independently for each subject by sampling uniformly from the integers 10 through 20. Let *x*_*i*1_ denote the binary covariate and *x*_*i*2_ denote the continuous covariate for subject *i*.

### 3.2. Shifted-Poisson Dwell-Time Simulations

Under the shifted-Poisson setting, subject- and state-specific dwell times were generated according to

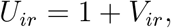

where

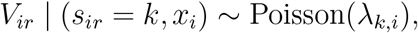

with

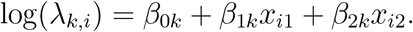

Consequently,

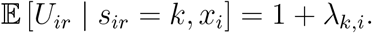

The true state-specific regression coefficients used for the shifted-Poisson simulations were

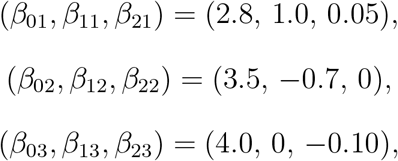

and

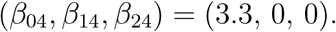

This setup was chosen so that the binary and continuous covariates influenced dwell times in some, but not all, states, thereby allowing assessment of both signal recovery and null-effect estimation. Under this parameterization, the regression coefficients describe covariate effects on *λ*_*k,i*_, which equals the expected dwell time minus one.

### 3.3. Gamma Dwell-Time Simulations

Under the Gamma setting, subject- and state-specific dwell times were generated from a Gamma distribution with a state-specific shape parameter and a covariate-dependent mean. Specifically,

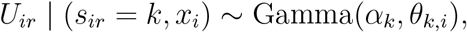

where *α*_*k*_ = 3 for all *k* = 1, …, 4. The covariate-dependent mean was defined as

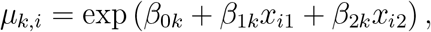

and the scale parameter was set to

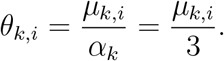

Thus,

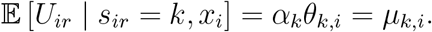

The true state-specific regression coefficients used for the Gamma simulations were

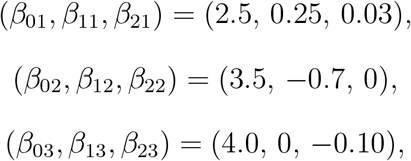

and

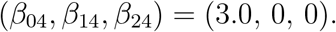

As in the shifted-Poisson setting, this design allowed covariate effects to vary across states, with some coefficients set to zero to assess estimation under the null. Under the Gamma mean–shape parameterization, the regression coefficients describe multiplicative covariate effects on the expected dwell time.

### 3.4. Generation of fMRI-Like Time Series

After generating the latent state sequences and corresponding dwell times, we simulated fMRI-like multivariate time series using the SimTB toolbox (Erhardt et al. (2012)). For each subject, we generated *p* = 10 component time courses over *T* = 500 time points with repetition time TR = 2 seconds. The simulated time series were constructed to reflect four distinct connectivity states, each characterized by a different module structure across the 10 components (see Figure 1).

**Figure 1:**
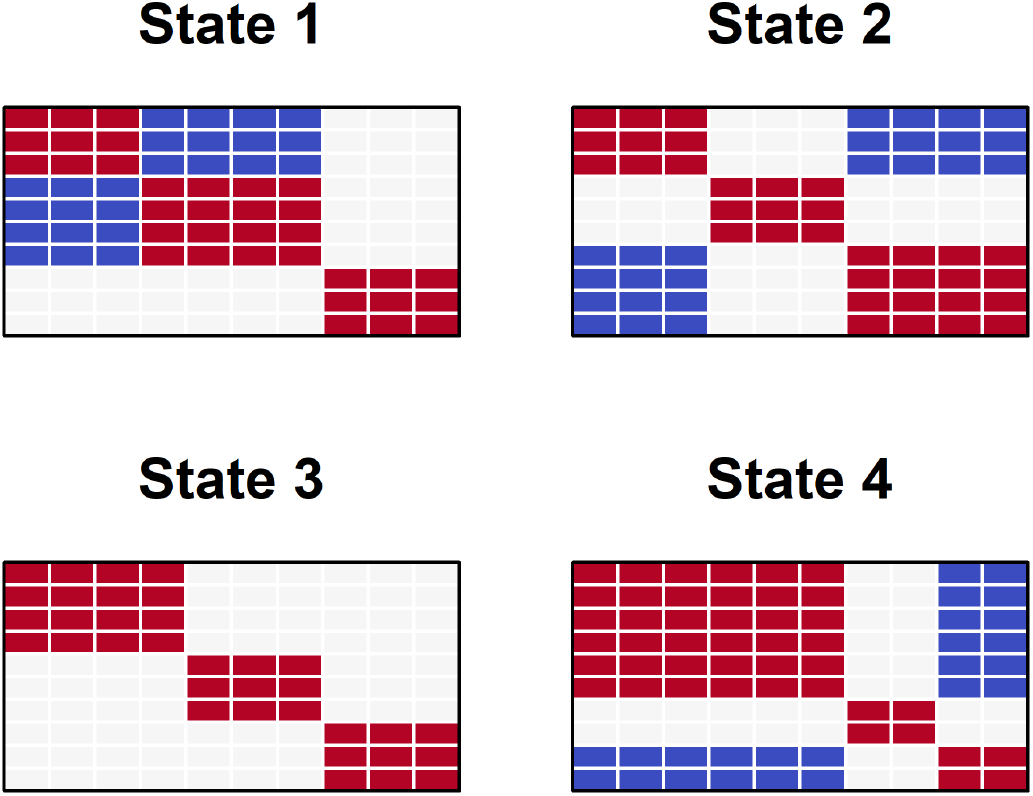
Connectivity structure of the four simulated network states used in the simulation study. Each panel shows the connectivity pattern among the 10 nodes for a given state. Red indicates positive connectivity between nodes, blue indicates negative connectivity, and white indicates no connectivity relationship. These state-specific structures were used within the SimTB framework to generate multivariate fMRI-like time series whose connectivity patterns change according to the simulated latent state sequence.

Within each state, components were assigned to modules through a prespecified state-dependent module membership matrix. Components belonging to the same module shared event-related activity, while components assigned opposite signs within a module exhibited negatively related activity. In addition to these state-dependent shared events, component-specific random events were added to induce subject-level and component-level variability. State-specific events occurred with probability 0.5, and unique component-level events also occurred with probability 0.5, with unique event amplitudes scaled to 0.5 relative to the shared module events.

For each subject, the simulated latent state sequence determined which connectivity state was active at each time point. Event time courses were then generated according to the active state’s module structure and convolved with a canonical hemodynamic response function to produce fMRI-like signals. Gaussian noise was subsequently added to each component time series. This procedure yielded multivariate time series whose dependence structure changed over time according to the simulated latent state process, thereby mimicking the key features of dynamic functional connectivity data.

For computational tractability, simulated networks contained *p* = 10 nodes. In real neuroimaging applications, the number of network nodes may be substantially larger (e.g., 50–200 regions of interest). The primary objective of the simulation study was to evaluate the ability of the proposed HSMM framework to recover covariate effects in the state dwell-time distributions rather than to study high-dimensional covariance estimation. Using a modest network size allowed us to isolate the performance of the dwell-time regression component while maintaining realistic state-dependent connectivity patterns generated through the SimTB toolbox. Because the dwell-time parameters are estimated through the latent state process rather than directly from the dimensionality of the observed network, increasing the number of nodes would primarily increase computational cost without substantially altering the dwell-time estimation problem.

### 3.5. Model Fitting and Evaluation

For each simulated dataset, we fit the proposed covariate-dependent HSMM using the correct number of latent states (*K* = 4). In the shifted-Poisson simulation setting, dwell times were modeled using the shifted-Poisson regression formulation, while in the Gamma simulation setting, dwell times were modeled using the Gamma mean–shape regression formulation. In both cases, the primary quantities of interest were the state-specific regression coefficients linking the binary and continuous covariates to the dwell-time component of the model.

Model performance was evaluated by comparing the estimated regression coefficients with their true values across simulation replicates. For each state and each regression coefficient, we computed the average estimated coefficient and the empirical standard deviation across simulated datasets. These summaries provide an assessment of both bias and variability in the estimated dwell-time regression parameters.

## 4. UK Biobank Application

To illustrate the proposed covariate-dependent HSMM framework in a real neuroimaging dataset, we applied the model to resting-state fMRI data from the UK Biobank imaging study. The UK Biobank is a large-scale biomedical database containing extensive imaging and health-related data from participants across the United Kingdom (Sudlow et al. (2015)).

### 4.1. Network Definition

Functional brain networks were defined using the group independent component analysis (ICA) decomposition provided by the UK Biobank imaging pipeline. Specifically, we used the 100-component group-ICA decom-position and retained the subset of components classified as “good” signal components by the UK Biobank quality-control procedure. Following UK Biobank guidelines, this resulted in 55 ICA components, which were used as the nodes of the network (see the UK Biobank imaging documentation: https://www.fmrib.ox.ac.uk/ukbiobank/). The associated component time series served as the multivariate observations for the dynamic connectivity analysis.

To facilitate interpretation of the connectivity structure, the retained ICA components were further grouped into canonical large-scale functional sub-networks based on prior UK Biobank work. Specifically, the components were organized into the anterior and posterior default mode networks (DMNa and DMNp), somatomotor network (SMN), subcortical and cerebellar network (Sub&Cereb), temporoparietal network (TempPar), ventral attention network (VAN), and visual network (VIS). These groupings were used to characterize within- and between-network connectivity patterns in the identified states (Jiang et al. (2022)).

### 4.2. Study Sample

A subset of 1,000 participants was selected from the UK Biobank imaging cohort. For each participant, the resting-state fMRI data were represented as a multivariate time series consisting of the 55 ICA component signals observed at 490 time points. These time series were analyzed using the HSMM framework described above.

### 4.3. Model Specification

We fit the proposed hidden semi-Markov model with *K* = 12 latent brain states. The use of 12 states was consistent with prior work in similarly sized Human Connectome Project datasets, in which models with 12 states have been used to characterize large-scale brain dynamics (Vidaurre et al., 2017; Shappell et al., 2019). Each state was characterized by a multivariate Gaussian emission distribution describing the joint activation pattern across the 55 ICA components. The state-specific covariance matrices represented the functional connectivity structure among the network nodes.

To model temporal persistence within states, dwell times were represented using the Gamma mean–shape specification described in Section 2. The mean dwell time was modeled as a function of subject-level covariates through a log-linear regression model, while the Gamma shape parameter was estimated separately for each state.

The dwell-time regression design matrix included an intercept, sex, and age. Sex was entered as a binary covariate, while age was entered as a continuous covariate. These covariates were allowed to affect the expected dwell time separately in each state, while the emission distributions and transition probability matrix remained common across participants.

A total of *B* = 500 subject-level bootstrap resamples were requested. Standard errors for the state-specific dwell-time regression coefficients were computed from the usable bootstrap replicates according to the procedure described in Section 2.6. Two-sided *p*-values were obtained from Wald statistics using a standard normal reference distribution. The number of bootstrap resamples was selected to balance computational feasibility with the Monte Carlo precision of the resulting uncertainty estimates.

## 5. Results

### 5.1. Simulation Results

We evaluated the performance of the proposed covariate-dependent HSMM by examining how accurately the model recovered the state-specific regression coefficients governing the dwell-time distributions. For each simulation scenario, model parameters were estimated for 100 independently generated datasets. Estimation performance was summarized using the mean estimated coefficient and the empirical standard deviation across simulation replicates.

#### 5.1.1. Shifted-Poisson Dwell-Time Simulations

Under the shifted-Poisson data-generating mechanism, the proposed HSMM closely recovered the true regression coefficients governing the dwell-time distributions across all four states. Table 1 summarizes the true parameter values and the average estimated coefficients across simulation replicates.

**Table 1:** Recovery of dwell-time regression coefficients under the shifted-Poisson simulation setting. Values shown are the mean estimated coefficients across 100 simulation replicates, with empirical standard deviations in parentheses.

|  | Intercept |  | Binary Covariate |  | Continuous Covariate |  |
|---|---|---|---|---|---|---|
| State | True | Estimate | True | Estimate | True | Estimate |
| State 1 | 2.8 | 2.792 (0.011) | 1.0 | 1.003 (0.004) | 0.05 | 0.050 (0.001) |
| State 2 | 3.5 | 3.495 (0.020) | -0.7 | -0.717 (0.010) | 0 | 0.000 (0.001) |
| State 3 | 4.0 | 3.987 (0.024) | 0 | 0.004 (0.011) | -0.10 | -0.100 (0.002) |
| State 4 | 3.3 | 3.297 (0.014) | 0 | -0.003 (0.006) | 0 | 0.000 (0.001) |

For states in which covariate effects were present, the estimated coefficients were close to their true values and had relatively small empirical standard deviations. For example, in State 1, the true intercept, binary covariate coefficient, and continuous covariate coefficient were 2.8, 1.0, and 0.05, respectively, while the corresponding average estimates were 2.792 (0.011), 1.003 (0.004), and 0.050 (0.001). Similarly, in State 2, the model recovered the negative binary covariate effect, with a true coefficient of −0.7 and an average estimate of −0.717 (0.010).

In states where covariate effects were absent, the estimated coefficients were centered near zero. For example, the binary covariate coefficient in State 3 had a true value of 0 and an average estimate of 0.004 (0.011), while the continuous covariate coefficient in State 4 had a true value of 0 and an average estimate of 0.000 (0.001). These results indicate that the model recovered both nonzero and null covariate effects under the shifted-Poisson specification.

Under this parameterization, the regression coefficients describe covariate effects on *λ*_*k,i*_, where

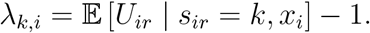

Thus, the coefficients represent multiplicative effects on expected dwell time minus one rather than on the complete expected dwell time.

#### 5.1.2. Gamma Dwell-Time Simulations

A similar pattern was observed when dwell times were generated under the Gamma mean–shape specification. Table 2 presents the true and estimated regression coefficients under the Gamma simulation scenario. Across the four states, the proposed HSMM generally produced estimates close to the true parameter values, although variability was greater than under the shifted-Poisson specification and modest attenuation was observed for several coefficients.

**Table 2:** Recovery of dwell-time regression coefficients under the Gamma simulation setting. Values shown are the mean estimated coefficients across 100 simulation replicates, with empirical standard deviations in parentheses.

|  | Intercept |  | Binary Covariate |  | Continuous Covariate |  |
|---|---|---|---|---|---|---|
| State | True | Estimate | True | Estimate | True | Estimate |
| State 1 | 2.5 | 2.496 (0.054) | 0.25 | 0.234 (0.027) | 0.03 | 0.029 (0.003) |
| State 2 | 3.5 | 3.327 (0.061) | -0.7 | -0.656 (0.022) | 0 | 0.001 (0.004) |
| State 3 | 4.0 | 3.846 (0.050) | 0 | 0.002 (0.021) | -0.10 | -0.088 (0.003) |
| State 4 | 3.0 | 2.953 (0.053) | 0 | -0.001 (0.024) | 0 | 0.003 (0.003) |

For example, in State 1, the true intercept, binary covariate coefficient, and continuous covariate coefficient were 2.5, 0.25, and 0.03, respectively, while the corresponding average estimates were 2.496 (0.054), 0.234 (0.027), and 0.029 (0.003). In State 2, the model recovered the negative association between the binary covariate and mean dwell time, with a true coefficient of −0.7 and an average estimate of −0.656 (0.022). The State 2 intercept was also estimated somewhat below its true value, with an average estimate of 3.327 compared with the true value of 3.5.

Coefficients corresponding to null covariate effects were estimated close to zero. For example, the binary covariate coefficient in State 3 had a true value of 0 and an average estimate of 0.002 (0.021), while the binary covariate coefficient in State 4 had a true value of 0 and an average estimate of −0.001 (0.024).

Under the Gamma mean–shape parameterization, the regression coefficients act directly on the conditional mean dwell time through

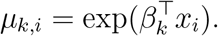

Consequently, exponentiated covariate coefficients represent multiplicative effects on expected dwell time.

Overall, the simulation results indicate that the proposed covariate-dependent HSMM recovered the state-specific dwell-time regression coefficients under both the shifted-Poisson and Gamma specifications. Recovery was particularly close under the shifted-Poisson setting. Under the Gamma setting, estimates remained generally close to their true values, although several coefficients exhibited modest attenuation and somewhat greater empirical variability.

### 5.2. UK Biobank Application

We next applied the proposed covariate-dependent HSMM to resting-state fMRI data from the UK Biobank imaging study to examine whether subject-level covariates were associated with differences in the temporal persistence of functional brain states.

#### 5.2.1. Sample Characteristics

The analysis included 1,000 participants from the UK Biobank imaging cohort. Table 3 summarizes the demographic characteristics of the study sample. The mean age of participants was 54.84 years (SD = 7.19), with ages ranging from 40 to 69 years. The sample consisted of 456 males and 544 females.

**Table 3:** Demographic characteristics of the UK Biobank sample used in the analysis.

| Characteristic | Value |
|---|---|
| Sample size (*N*) | 1,000 |
| Male | 456 |
| Female | 544 |
| Age, mean (SD) | 54.84 (7.19) |
| Age range | 40–69 |

#### 5.2.2. Covariate Associations with Mean State Dwell Time

Table 4 presents the estimated state-specific regression coefficients for sex and age. Under the Gamma mean–shape parameterization, positive coefficients indicate higher expected dwell times as the corresponding covariate increases, whereas negative coefficients indicate lower expected dwell times. Exponentiated coefficients represent multiplicative effects on expected dwell time.

**Table 4:**
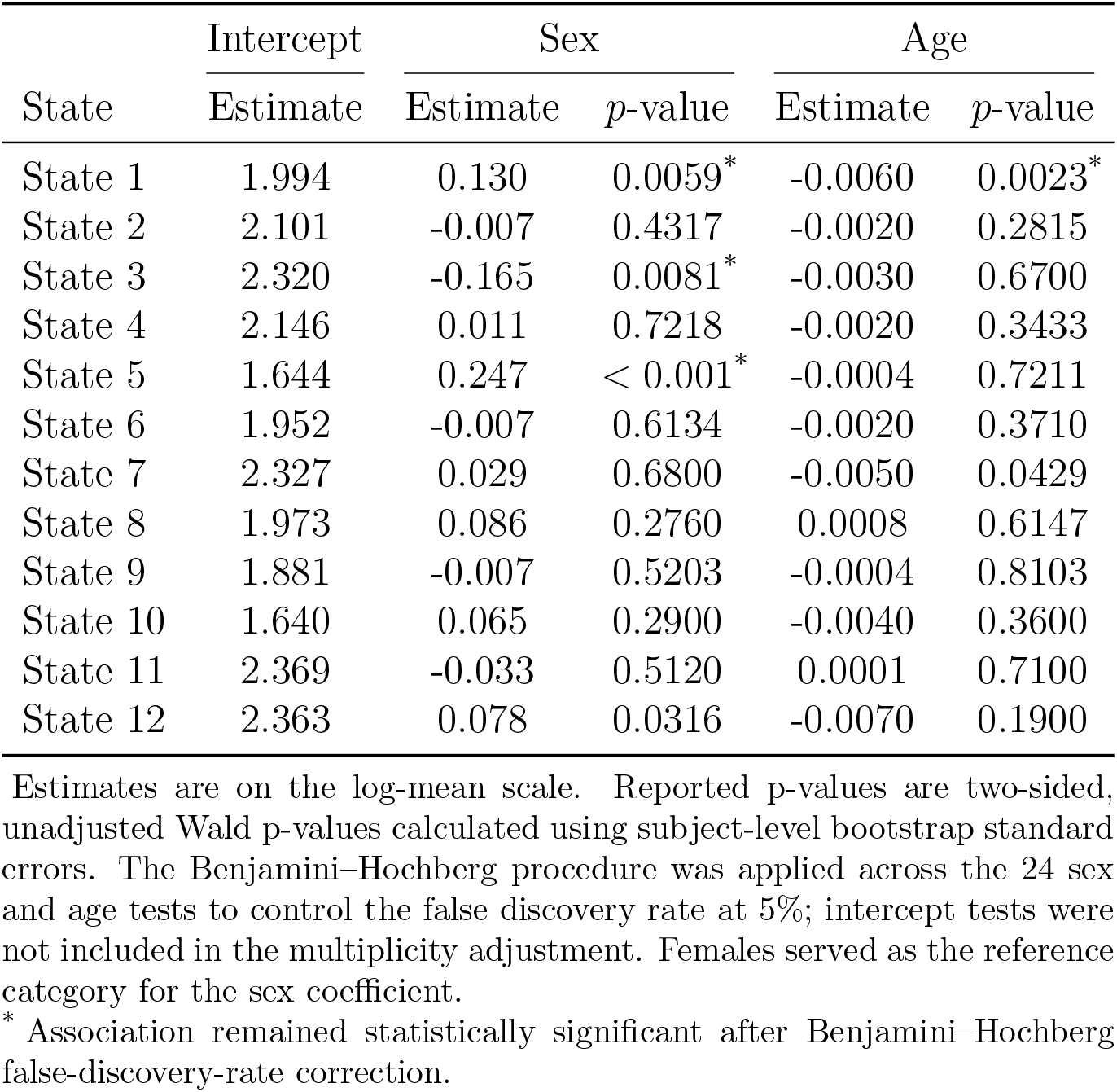
Estimated regression coefficients for associations of sex and age with mean dwell time across the 12 latent brain states.

|  | Intercept | Sex |  | Age |  |
|---|---|---|---|---|---|
| State | Estimate | Estimate | *p*-value | Estimate | *p*-value |
| State 1 | 1.994 | 0.130 | 0.0059<sup>*</sup> | -0.0060 | 0.0023<sup>*</sup> |
| State 2 | 2.101 | -0.007 | 0.4317 | -0.0020 | 0.2815 |
| State 3 | 2.320 | -0.165 | 0.0081<sup>*</sup> | -0.0030 | 0.6700 |
| State 4 | 2.146 | 0.011 | 0.7218 | -0.0020 | 0.3433 |
| State 5 | 1.644 | 0.247 | < 0.001<sup>*</sup> | -0.0004 | 0.7211 |
| State 6 | 1.952 | -0.007 | 0.6134 | -0.0020 | 0.3710 |
| State 7 | 2.327 | 0.029 | 0.6800 | -0.0050 | 0.0429 |
| State 8 | 1.973 | 0.086 | 0.2760 | 0.0008 | 0.6147 |
| State 9 | 1.881 | -0.007 | 0.5203 | -0.0004 | 0.8103 |
| State 10 | 1.640 | 0.065 | 0.2900 | -0.0040 | 0.3600 |
| State 11 | 2.369 | -0.033 | 0.5120 | 0.0001 | 0.7100 |
| State 12 | 2.363 | 0.078 | 0.0316 | -0.0070 | 0.1900 |

Estimates are on the log-mean scale. Reported p-values are two-sided, unadjusted Wald p-values calculated using subject-level bootstrap standard errors. The Benjamini–Hochberg procedure was applied across the 24 sex and age tests to control the false discovery rate at 5%; intercept tests were not included in the multiplicity adjustment. Females served as the reference category for the sex coefficient.
<sup>\*</sup> Association remained statistically significant after Benjamini–Hochberg false-discovery-rate correction.

At the nominal, unadjusted 0.05 significance level, sex was associated with mean dwell time in States 1, 3, 5, and 12. Under the sex coding used in the analysis, positive coefficients in States 1, 5, and 12 indicated longer expected dwell times for males relative to females, whereas the negative coefficient in State 3 indicated shorter expected dwell times for males relative to females. Age was nominally associated with mean dwell time in States 1 and 7, with negative coefficients indicating shorter expected dwell times with increasing age.

To account for multiple testing, the p-values for the 24 sex and age tests across the 12 states were adjusted using the Benjamini–Hochberg procedure to control the false discovery rate at 5%. Following this correction, the associations between sex and mean dwell time in States 1, 3, and 5 remained statistically significant, as did the association between age and mean dwell time in State 1. The corresponding false-discovery-rate-adjusted p-values were 0.0472 for the sex effect in State 1, 0.0486 for the sex effect in State 3, 0.0210 for the sex effect in State 5, and 0.0276 for the age effect in State 1. The sex association in State 12 and the age association in State 7 did not remain significant following false-discovery-rate correction, with adjusted p-values of 0.1517 and 0.1716, respectively. To aid interpretation of the effect magnitudes, the regression coefficients were exponentiated. Among the associations that remained significant after false-discovery-rate correction, male sex was associated with an approximately 13.9% longer expected mean dwell time in State 1, a 15.2% shorter expected mean dwell time in State 3, and a 28.0% longer expected mean dwell time in State 5 relative to female sex, holding age constant. In State 1, each 10-year increase in age was associated with an approximately 5.8% shorter expected mean dwell time, holding sex constant.

These results indicate that associations between demographic characteristics and temporal persistence differed across latent brain states. Of the 500 requested subject-level bootstrap resamples, 491 satisfied the convergence criterion and were used for statistical inference.

#### 5.2.3. Estimated Dwell-Time Distributions

To illustrate the estimated covariate associations, we examined the fitted Gamma density curves at selected sex and age values. Figure 2 displays these fitted density curves for each of the 12 latent states.

**Figure 2:**
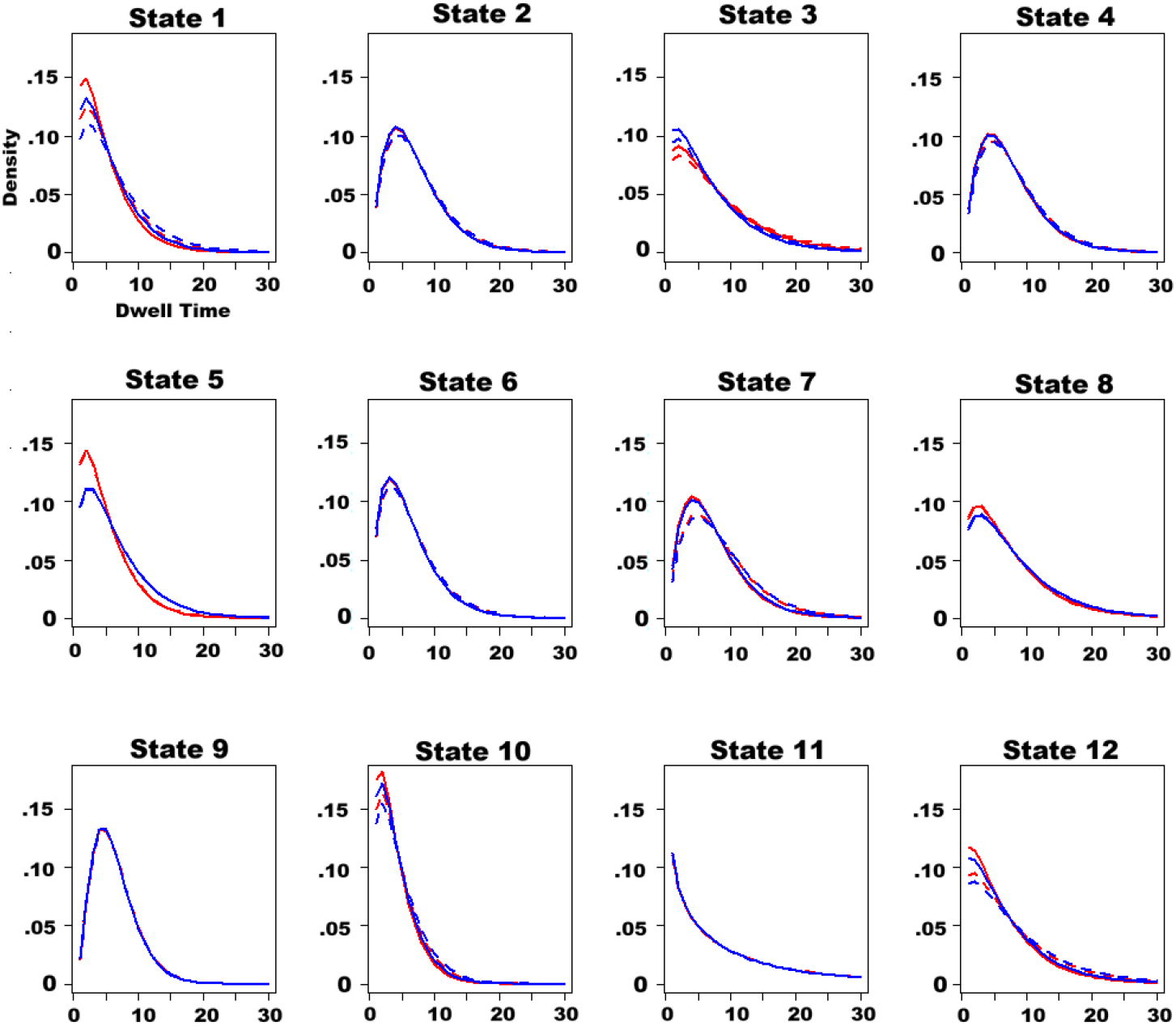
Fitted Gamma density curves for the 12 latent brain states identified in the UK Biobank analysis. Curves are stratified by sex (blue = male, red = female) and age (solid = 69 years, dashed = 40 years). Differences between curves arise from covariate-dependent changes in the conditional mean and scale of the Gamma distribution; the state-specific shape parameter is common across covariate profiles within each state.

Because sex and age entered the model through the conditional mean, differences between curves reflect covariate-dependent changes in the mean and scale of the Gamma distribution. The state-specific Gamma shape parameter remained common across covariate profiles within each state.

In states with nominal sex associations, the fitted density curves showed differences consistent with the signs of the regression coefficients. In States 1, 5, and 12, the fitted curves for males were shifted toward longer dwell durations relative to those for females. State 3 showed the opposite pattern, consistent with its negative sex coefficient.

The nominal age associations in States 1 and 7 were reflected in modest shifts toward shorter dwell durations at older ages. In most other states, the fitted curves showed substantial overlap across the displayed sex and age profiles, consistent with the smaller estimated covariate effects.

#### 5.2.4. Connectivity Structure of States with Nominal Covariate Associations

To characterize the connectivity patterns associated with the states showing nominal covariate associations, we examined the state-specific correlation matrices derived from the fitted Gaussian emission covariance matrices (Figure 3).

**Figure 3:**
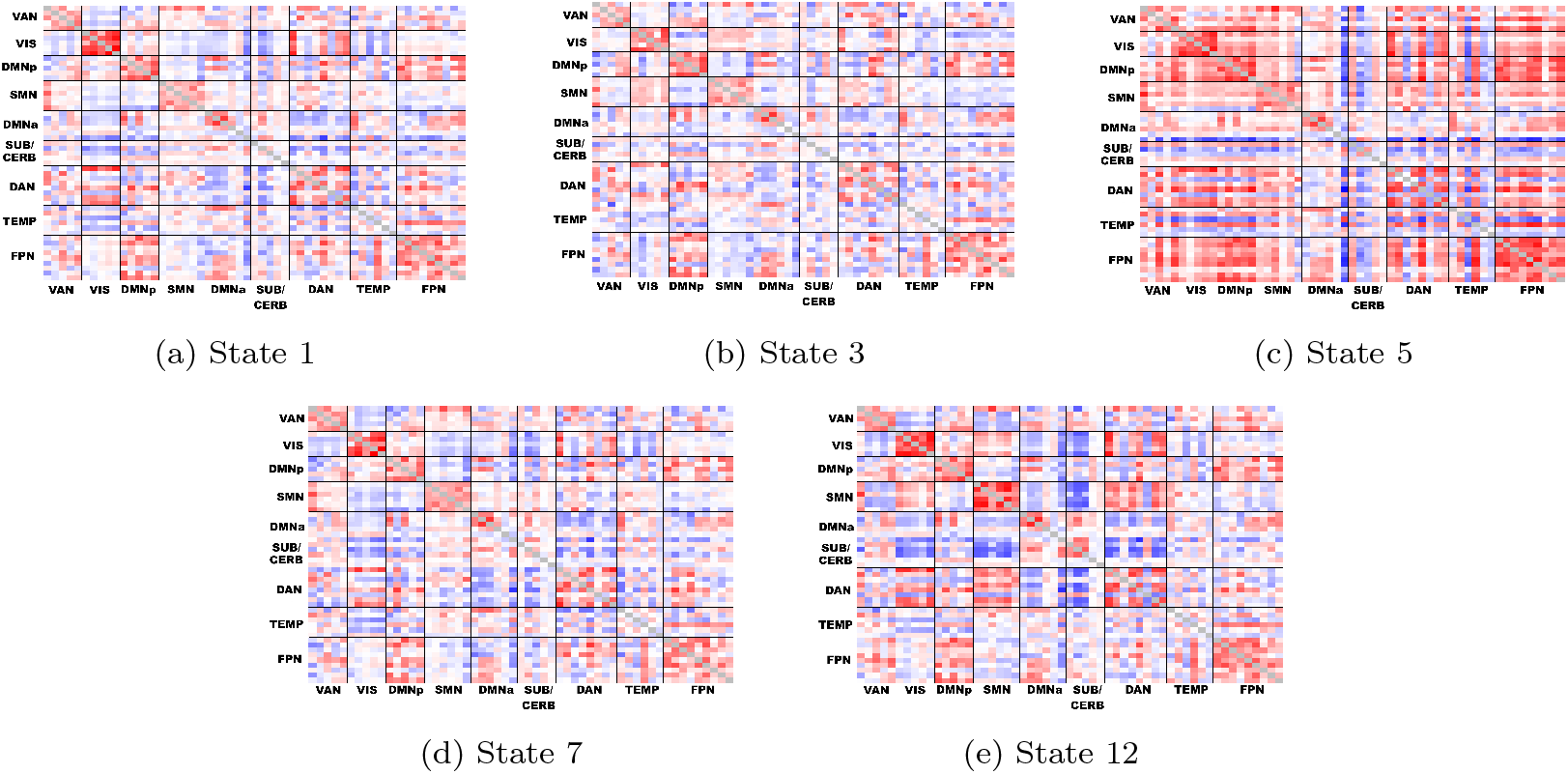
State-specific functional connectivity correlation matrices for the states exhibiting nominal, unadjusted associations between the covariates and mean dwell time. Regions are ordered by subnetwork, with independent components grouped into the anterior and posterior default mode networks (DMNa and DMNp), somatomotor network (SMN), subcortical and cerebellar network (Sub&Cereb), temporoparietal network (TempPar), ventral attention network (VAN), and visual network (VIS). A common color scale is used across panels, with blue indicating negative correlations and red indicating positive correlations.

Among the displayed states, State 5 exhibited the greatest overall contrast, with relatively strong positive and negative correlations distributed across multiple subnetworks. State 12 also showed a comparatively distinct pattern, including pronounced negative correlations between subcortical and cerebellar components and several cortical subnetworks. States 1, 3, and 7 exhibited more moderate connectivity patterns, with differences primarily in the magnitude and distribution of within- and between-network correlations.

These results illustrate that states with covariate-dependent differences in temporal persistence can correspond to distinct functional connectivity configurations. However, the connectivity descriptions are qualitative and are intended to aid interpretation rather than constitute formal comparisons between state-specific matrices.

## 6. Discussion

In this work, we developed a covariate-dependent extension of the hidden semi-Markov model (HSMM) that allows time-invariant, subject-level variables to influence the state-specific dwell-time distributions of latent brain states. Standard multi-subject HSMM formulations commonly assume that dwell-time behavior is shared across individuals, limiting the ability to assess how subject characteristics relate to temporal persistence within particular states. By incorporating state-specific fixed effects into the dwell-time component, the proposed framework provides an interpretable approach for studying how expected state durations vary across individuals and populations.

The simulation results indicate that the proposed model can recover state-specific covariate effects under both shifted-Poisson and Gamma dwell-time specifications. Recovery was particularly close under the shifted-Poisson specification, with estimated regression coefficients centered near their true values and relatively small empirical standard deviations. Under the Gamma specification, the estimated coefficients also generally approximated their true values, although several coefficients exhibited modest attenuation and somewhat greater variability. Together, these findings support the ability of the EM-based estimation procedure to identify both nonzero and null covariate effects under the simulation conditions considered.

In the UK Biobank application, sex and age were associated with expected dwell time in a subset of states, and the estimated associations were state-specific. At the nominal, unadjusted 0.05 level, sex was associated with mean dwell time in States 1, 3, 5, and 12, while age was associated with mean dwell time in States 1 and 7. After applying the Benjamini–Hochberg procedure across the 24 sex and age tests to control the false discovery rate at 5%, the sex associations in States 1, 3, and 5 and the age association in State 1 remained statistically significant. The sex association in State 12 and the age association in State 7 did not remain significant after correction. These results suggest that demographic associations with temporal persistence are localized to particular functional configurations rather than uniformly expressed across all states.

The fitted dwell-time distributions further illustrated the state-specific nature of these associations. In states with estimated covariate effects, the curves differed through covariate-dependent changes in the conditional Gamma mean and scale, while the state-specific shape parameter remained common across covariate profiles. Many states showed substantial overlap across the displayed sex and age profiles, consistent with the absence of strong covariate associations in those states.

The connectivity matrices provided additional qualitative context for the states exhibiting nominal covariate associations. Among the displayed states, State 5 exhibited relatively strong positive and negative correlations across multiple subnetworks, whereas State 12 showed pronounced negative correlations between subcortical and cerebellar components and several cortical subnetworks. States 1, 3, and 7 exhibited more moderate differences in the magnitude and distribution of within- and between-network correlations. These descriptions are intended to aid interpretation of the identified states rather than constitute formal statistical comparisons between their connectivity matrices. Accordingly, the results do not establish that particular connectivity features cause the observed differences in dwell time.

Prior work has associated the rate of switching between network configurations with cognitive flexibility and performance (Pedersen et al., 2018; Chen et al., 2025). The present findings complement this work by demonstrating that expected dwell time can vary systematically with subject-level characteristics and that these associations may differ across functional connectivity states. This emphasizes the value of examining not only which network configurations are expressed, but also the temporal persistence of each configuration.

The age-related findings provide an example of this state-specific temporal variation. The negative age coefficient in State 1 remained significant after false-discovery-rate correction, indicating that increasing age was associated with shorter expected visits to this particular state. A similar negative coefficient was observed in State 7 at the nominal level, although this association did not remain significant after correction. Previous studies have reported age-related differences in large-scale brain organization (Betzel, 2014; Farras-Permanyer et al., 2019). The present results suggest that such differences may also extend to the temporal persistence of selected network configurations. However, shorter dwell time in an individual state should not be interpreted as evidence of uniformly faster switching across the complete state process.

Sex was also associated with mean dwell time in a subset of states. The associations in States 1, 3, and 5 remained significant after false-discovery-rate correction, while the association in State 12 was nominally significant but did not survive correction. Prior studies have identified sex-related differences in functional network organization and connectivity strength (Scheinost et al., 2015; Zhang et al., 2018). The present findings suggest that sex-related differences may also be reflected in the expected duration of selected functional brain states. Because the observed associations were not consistent across all states, they should be interpreted as state-specific differences in temporal persistence rather than a global difference in brain-state stability.

Taken together, these findings highlight the importance of jointly characterizing the spatial and temporal properties of functional brain networks. Traditional functional connectivity analyses frequently rely on static summaries, while dynamic analyses may focus primarily on identifying states and transitions between them. The proposed framework complements these approaches by explicitly modeling dwell-time behavior and allowing its conditional mean to vary with subject-level covariates.

Several limitations should be considered. First, the proposed model assumes that the emission distributions, transition probabilities, and initialstate distribution are shared across subjects, with only the dwell-time component allowed to vary with covariates. The model also treats subject-level covariates as time-invariant fixed effects and does not incorporate subjectspecific random effects. Extending the framework to include random effects or to allow the emission and transition parameters to depend on covariates would be valuable, particularly for repeated-measures and longitudinal studies. For example, covariate-dependent transition probabilities could be incorporated within the EM framework by using the posterior expected transition counts from the E-step as fractional counts in origin-state-specific multinomial logistic regressions in the M-step. Although conceptually straight-forward, this extension would substantially increase the number of parameters. With *K* states and *q* covariate-design variables, a baseline-category parameterization requires *K*(*K* − 2)*q* transition-regression coefficients when self-transitions are excluded. The number of parameters therefore grows approximately quadratically with the number of states, potentially increasing computational burden and reducing estimation stability when some origin– destination transitions are infrequent. Regularized or structured models for covariate-dependent transition probabilities represent an important direction for future work.

Second, specification of the number of latent states remains an important consideration. In the simulation study, the model was fit using the true number of states, allowing the evaluation to focus on regression-parameter recovery. In the UK Biobank application, *K* = 12 was selected based on prior work using similarly sized neuroimaging datasets, and we did not perform a formal sensitivity analysis across alternative values of *K*. The reported sex and age associations should therefore be interpreted as conditional on the selected 12-state representation. Alternative values of *K* may split, merge, or reorganize latent states and could consequently alter state correspondence and the estimated covariate effects. Future work should assess the robustness of covariate findings across a range of plausible state solutions, using state-alignment procedures to determine whether the directions and magnitudes of the identified associations are preserved across values of *K*.

Third, the Gamma implementation evaluates a continuous Gamma density at integer dwell durations and uses the resulting density values directly in the E-step. These values are not obtained by integrating the Gamma density over unit-width intervals and are not renormalized over the finite dwell-time grid. The Gamma specification should therefore be viewed as an integer-grid density approximation to the dwell-time contribution. In addition, because the Gamma and shifted-Poisson specifications use continuous density values and discrete probability masses, respectively, direct comparisons of their raw likelihoods or likelihood-based information criteria should be interpreted descriptively.

Fourth, the subject-level bootstrap is computationally intensive, particularly for large neuroimaging datasets and models containing many states. Only 500 bootstrap resamples were requested for the UK Biobank analysis, which may limit the Monte Carlo stability of the resulting standard errors, confidence intervals, and p-values. Bootstrap replicates may also fail to converge or produce unusable estimates, further reducing the effective number of replicates available for inference. Future applications would benefit from a larger number of resamples when computational resources permit. Moreover, although the simulation study evaluated recovery of the dwell-time regression coefficients, it did not directly assess the finite-sample calibration of the bootstrap standard errors, confidence intervals, or hypothesis tests. Future work should evaluate confidence-interval coverage, Type I error, and power under a broader range of sample sizes and model conditions.

Fifth, the UK Biobank analysis was observational and cross-sectional. The identified associations therefore should not be interpreted as causal effects of age or sex on brain-state persistence or as evidence of within-person changes over time. Furthermore, interpretation of latent states remains inherently data-driven, and linking the identified connectivity patterns and dwell-time differences to particular cognitive or behavioral processes will require external validation.

Despite these limitations, the proposed covariate-dependent HSMM provides a useful framework for studying individual differences in the temporal dynamics of brain networks. By enabling state-specific inference on expected dwell time, the approach offers a way to investigate how demographic, clinical, and biological characteristics relate not only to the expression of functional brain states, but also to the duration of visits to those states.

## Data and Code Availability

Data used in this study were obtained from the UK Biobank, which is a publicly available resource accessible through (https://www.ukbiobank.ac.uk). Access to individual-level data is subject to UK Biobank’s data access policies and requires appropriate approvals. Code implementing the proposed covariate-dependent HSMM is publicly available at https://github.com/rlyday/HSMM_Covariate_Model.

## Funding

This work was supported by the National Institute of Biomedical Imaging and Bioengineering (K25-EB032903-01).

## Declarations of Interest

All authors report no biomedical financial interests or potential conflicts of interest.

## Notes

### Competing Interest Statement

The authors have declared no competing interest.

